# fragilityindex: An R Package for Statistical Fragility Estimates in Biomedicine

**DOI:** 10.1101/562264

**Authors:** Kipp W. Johnson, Eli Rappaport, Khader Shameer, Benjamin S. Glicksberg, Joel T. Dudley

## Abstract

Poor reproducibility is a growing crisis in biomedical research. The fragility index was introduced as a convenient measure to estimate how fragile statistical results in clinical trials are to small perturbations in event outcome counts. There is currently no freely available R package to produce this calculation. Furthermore, the original definition of the method is applicable only to 2x2 contingency tables.

As such, we developed an R package to calculate fragility index. We have also extended the concept of a statistical fragility index to two of the most commonly used methods in clinical research, survival analysis via weighted log-rank tests and logistic regression, and implemented these technique sin this R package. We describe example applications of these methods to existing publically available datasets. This R package is freely available under the AGPL license on CRAN (https://cran.r-project.org/web/packages/fragilityindex/index.html). The most recent versions may be downloaded and installed via Github (https://github.com/kippjohnson/fragilityindex).

## Introduction

Awareness of the problem of poor reproducibility in biomedical science has grown in the past several years. There have been numerous attempts to explain this phenomenon, such as an over-reliance on null-hypothesis significance testing (1, 2), financial conflicts of interest, or simply prevailing biases in hypotheses tested (3). In 2014, Walsh et al. showed that the statistical significance of randomized clinical trials in medical research are often fragile, or sensitive, to small perturbations in patient outcomes with a simple statistic called the fragility index (4). The fragility index was originally defined for use in 2x2 contingency tables with dichotomous outcomes. In the case of clinical trials, fragility index is defined as the minimum number of patients it would take to make a statistically significant result non-significant if the patients’ outcomes were reversed. Despite being quite simple, this is a relevant statistic that can facilitate better interpretation of results from clinical trials. For example, Ridgeon et al. found that 40% of trials in critical care medicine had a fragility index of 1 or less (5), which is of large concern in the medical field.

This statistic has since been embraced by physicians in a number of clinical fields as an easy-to-interpret measure of the robustness of clinical trials (5, 6, 7). However, the adoption of this metric in routine statistical analyses has been limited due to the unavailability of a user-friendly statistical package. Here, we present an R package (8) “fragilityindex” which can perform fragility index estimations and has tools which extend the concept of the fragility index to two other statistical techniques often employed in medical research: survival analysis with weighted log-rank tests and logistic regression.

## Materials and Methods

### Fragility Index for contingency tables

Fragility index calculation is most often associated with contingency tables and there are many studies that portray its utility. For example, consider the following scenario: in a clinical trial, there are two groups of patients. In group one, 15/40 patients have an adverse event. Within group two, 6/40 patients have the adverse event. The corresponding exact test P-value for this situation is 0.041. However, if a single additional patient in group two experienced the adverse event, we obtain a new p-value of 0.078. This result is no longer reportable as “significant,” and the fragility index for this trial is 1. We consider this to be a fragile clinical trial result, since a single swapped outcome would change the conclusion of the study. In our package, we can conduct this test as follows:

> fragility.index(15, 6, 40, 40, verbose=TRUE)

fragility.index p.value

1. 0 0.041
2. 1 0.078

To further illustrate the utility of our tool, we applied this analysis to the results from all clinical studies publically available on ClinicalTrials.gov (https://www.ctti-clinicaltrials.org/aact-databaseaact-database, March 27th, 2016 accession). Before the data can be read into R, “line feed” characters within variables need to be removed from the datasets, which can be accomplished with the following line of bash code (taken from the README file included within the data):

> tr -d ’\012’ < clinical_study_noclob.txt > clinical_study_noclob_nolf.txt

As many of these studies are not in a useable format for the current analysis, we filtered them based on criteria described in Figure 1. Briefly, we only considered clinical trials where there were significant results (defined as p<0.05) from Fisher Exact tests performed on 2x2 tables. There are several issues with formatting in the dataset that must be adjusted before the resulting values can be used in our tests. Furthermore, some entries have impossible results (i.e. non-count values) where perhaps the statistical test was mis-specified on ClinicalTrials.gov. Due to limitations in the dataset, aggregate results should thus be considered with these limitations in mind. In the manuscript, we highlight the application of this function to a particular clinical trial whose values we manually verified. We provide the complete R code to generate the final fragility indices from the clinical data in the Supplemental Materials.

**Figure 1:**
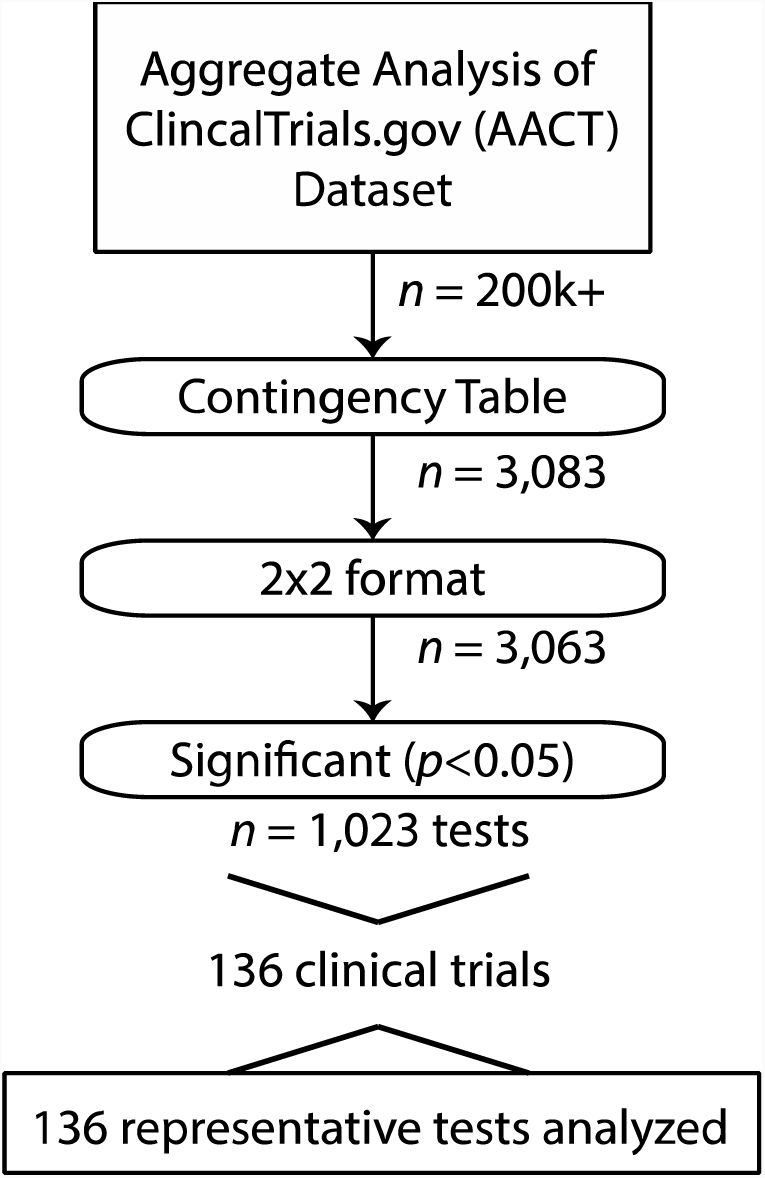
Workflow of filtering steps for available clinical trial data used in fragility index analysis for contingency tables. Data was acquired from ClinicalTrials.gov, specifically the publically available Aggregate Analysis of ClinicalTrials.gov (AACT) database.

### Fragility Index for survival analysis

To increase power, many clinical trials compare the outcomes of two groups studied longitudinally with survival analysis. We have developed a new technique to enable fragility index calculation in survival analysis utilizing the *survdiff()* function from the R package *“survival”* (9). A survival time series consists of paired outcomes (0/1) and exposure times for patients. To compute this fragility index analogously to the fragility index for 2x2 tables, we add a new individual to the input dataset with an exposure time equal to the average exposure time of all previous individuals. We indicate this individual died at the average exposure time. The individual’s covariate values, if applicable for the *survdiff()* function, are randomly sampled from another individual in the dataset. We then repeat the process for multiple iterations and average the outcome of each run into an aggregate, overall fragility index estimate for the dataset. This fragility index may thus be interpreted as the mean number of individuals it would take who died at the average exposure time of all individuals in the study to take a result from significant to non-significant.

To show how fragility index calculation can be extended to survival analyses, we performed our method on open-source TCGA, using a Colorectal Adenocarcinoma dataset with 276 patients (10). This dataset is comprised of patient survival rates along with their genetic profiles, particularly which genes have copy number alterations (CNA; amplifications or deletions). We then selected two genes, BCL2L1 and POFUT1, in which there were the most patients (35/276) with amplifications in both. The downloading and pre-processing steps are described fully in the supplementary material. Briefly, the TCGA dataset for colorectal adenocarcinoma (“coadread“) was downloaded from http://www.cbioportal.org/study?id=coadread_tcga_pub#summarystudy?id=coadread_tcga_pub#summary using the GUI option to “download data” on November 7th, 2016. From this file, we use the table entitled “data_clinical.txt” for survival information. For the CNA patient survival data, we used the GUI on the same website to check off genes BCL2L1 and POFUT1 and then used the GUI option to download this data, saving the file as “CNA_genes.txt“. We use patient identifiers from this file in the resulting analysis. Calculation of the survival analysis fragility index from these datasets can be performed with the R code located in the Supplementary Materials, using the R package “*survival*” for weighted log-rank tests.

### Fragility Index for Logistic Regression β-coefficients

We present a method to calculate logistic regression coefficient fragility, or how many events it would take to change a significant logistic regression coefficient to non-significant at the given confidence level. To do this, we replace responses (which should be binary events, 0/1) with the opposite event until the coefficient is non-significant. If the initial regression coefficient (β) is positive, we change a 1 event to a 0. If the regression coefficient is negative, we change a 0 event to a 1. We then count the number of times a replacement must occur to produce a non-significant result and use this amount as the fragility index. To account for variability in the response replacement, we then repeat this process a number of times and take the mean of all of the computed fragility indices as the representative fragility index. As with all multiple regression models, the issue of potential (multi)colinearity, or high correlation between variables, must be addressed before running the analysis. This is especially important as multicollinear variables may have a significant impact upon the “fragility” of the regression coefficients by making them unstable. Using existing packages, we must include this step as a check before running the fragility index for regression coefficients in order to most accurately interpret results.

## Results

### Fragility Index for contingency tables

We calculated the fragility index for a curated list of all available clinical trial studies from clinicaltrials.gov (https://www.ctti-clinicaltrials.org/aact-database, March 27th, 2016 accession). Of 3,083 Fisher’s exact test results, we excluded results that were not in a 2x2 format (n=19), were not statistically significant (p≥0.05, n= 2004), or had improper values (n=37), leaving 1,023 usable tests from 136 unique clinical trials (Figure 2). We randomly selected one test from each trial to calculate fragility index. We found five tests from five separate trails that have a fragility index of 1. For example, in a clinical trial (NCT00058019) evaluating the use of Ixabepilone (a chemotherapeutic agent) for non-Hodgkin’s lymphoma, one tested outcome was the response rate between chemosensitive and chemoresistant individuals. They found that Ixabepilone administered to chemosensitive individuals (14/39) had a better response rate (p=0.022) than chemoresistant individuals (0/12). We can apply the fragility index assessment on these outcomes:

> fragility.index(15, 6, 40, 40, verbose=TRUE)

fragility.index p.value

1. 0 0.041
2. 1 0.078

**Figure 2:**
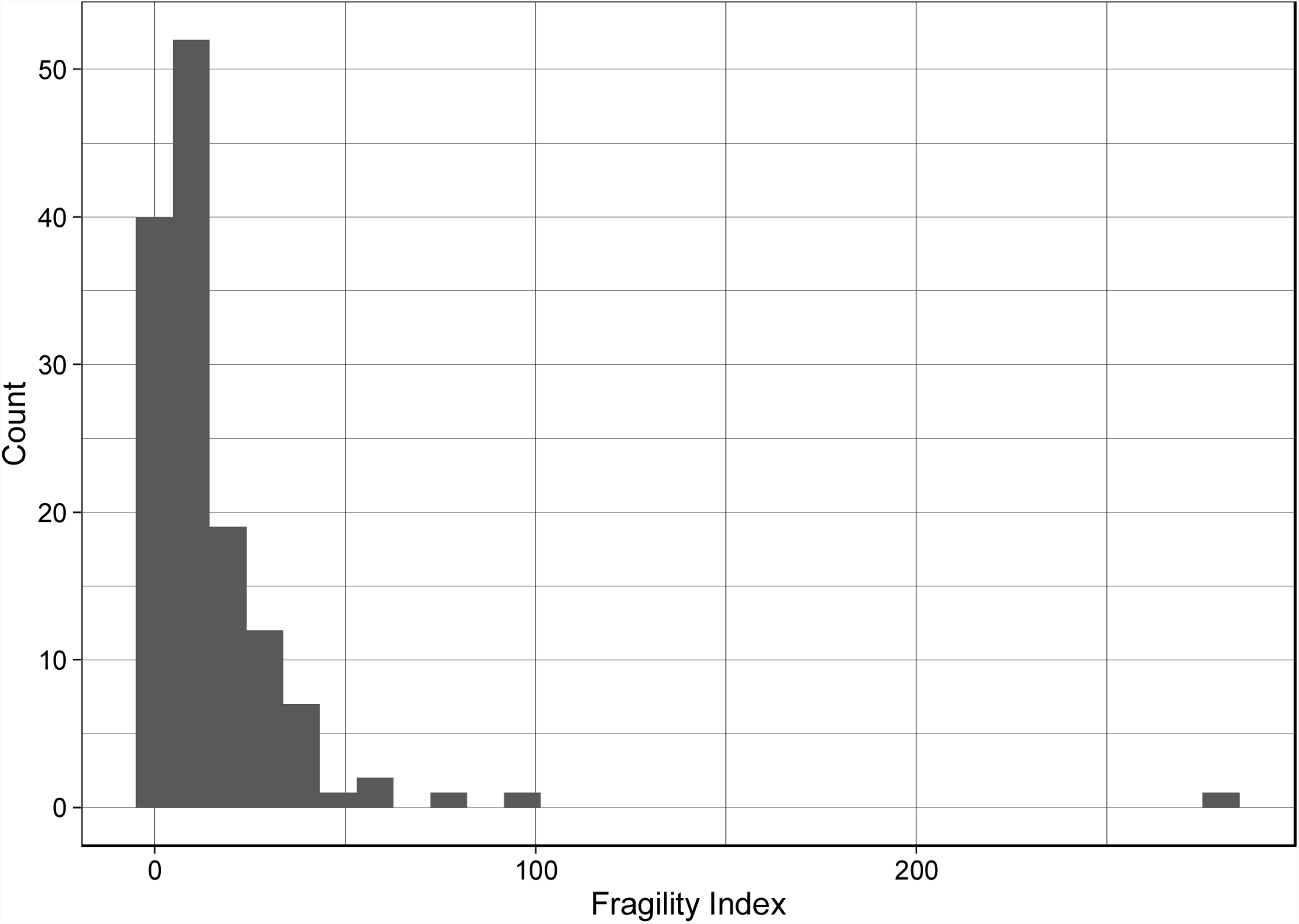
The distribution of fragility indices for the 136 tests used as representative examples from 136 distinct clinical trials.

Accordingly, a single addition in response outcome (i.e. a fragility index of 1) between the two groups makes the association not reportable as statistically significant (p=0.083). Reproducing or replicating the study to determine whether this effect was due simply to chance may be recommended. We provide full data acquisition steps and the fragility index distribution for all usable clinical trials in the Supplemental Materials.

### Fragility Index for Survival Analysis

We have applied this analysis to a Colorectal Adenocarcinoma dataset from TCGA (10) which assesses the effect of gene Copy Number Alteration on survival. Specifically, we calculated the fragility index for the significant association of amplifications of genes *BCL2L1* and *POFUT1* and survival (*p*=0.03) as follows (detailed steps to replicate analysis in Supplemental Materials):

~~~
survivalfragility(Surv(ronths, status) ∼ cna.status, data=infile,
   niter=100, progress.bar=TRUE)
 |++++++++++++++++++++++++++++++++++++++++++++++++++| 100% elapsed = 06s
[1] 1.36
~~~

Although the survival analysis is significant for amplification of these 2 genes, we find that it is relatively fragile as it is non significant by the addition of only 1-2 individuals who die at the mean exposure time.

### Fragility Index for Logistic Regression β-coefficients

To illustrate the utility of calculating a fragility index for logistic regression, we utilized data from UCI Machine Learning Repository (11), specifically a multivariate heart disease dataset (Cleveland site). This dataset is comprised of an outcome variable (heart disease status) and 13 predictor variables (full description can be found on their website). We provide the full R code to reproduce this example analysis in the Supplementary Materials.

~~~
# before running logisticfragility function, first test for (multi)colinearity
(see Supplementary Materials)
> logisticfragility(num ∼ age + sex +cp + trestbps + chol + fbs + restecg +
thalach + exang + oldpeak + slope + ca + thal, data = mydata, covariate=“all“,
niter=100, pro-gress.bar=FALSE)
# only significant associations displayed
  coefficient fragility.index
1 (Intercept)  17.21
3    sex  29.81
4     cp  35.73
5trestbps  8.06
9 thalach  4.99
10 exang   9.37
13    ca  69.69
14  thal 61.83
~~~

We find that the “ca” (# of major vessels colored by flourosopy) and “thal” (defect type) coefficients are the least fragile as heart disease predictors, while “thalach” (maximum heart rate achieved), “trestbps” (resting blood pressure), and “exang” (presence of exercised induced angina) are the most fragile. As previously indicated (6, 12), we do find a positive correlation between degree of significance and fragility (i.e. higher significance generally leads to more stable fragility scores) and, as such, both metrics should be considered when interpreting outcome associations.

## Discussion

We have developed the first R statistical package for the calculation of fragility index in multiple common settings, having extended and incorporated this technique to other commonly used statistical tests in the medical literature. Our extensions of the fragility index to survival analysis and logistic regression may prove to be helpful tools for researchers attempting to analyze clinical results. We hope that increased awareness of the fragility of some results will lead re-searchers to be more cognizant of the fragility of their claims. Ultimately, we hope this process will contribute to enhancing the reproducibility of the biomedical literature.

## Acknowledgements

We would like to thank Institute of Next Generation Healthcare, Mount Sinai Health System, New York, NY for infrastructure and support. We would also like to thank the authors of the open-source datasets used as examples for this study.

## References

1. Leggett NC, Thomas NA, Loetscher T, Nicholls ME. The life of p: “just significant” results are on the rise. Q J Exp Psychol (Hove) 2013;66: 2303–2309.

2. Glaser DN. The controversy of significance testing: misconceptions and alternatives. Am J Crit Care 1999;8: 291–296.

3. Ioannidis JP. Why most published research findings are false. PLoS Med 2005;2:e124.

4. Walsh M, Srinathan SK, McAuley DF, Mrkobrada M, Levine O, Ribic C, et al. The statistical significance of randomized controlled trial results is frequently fragile: a case for a Fragility Index. J Clin Epidemiol 2014;67: 622–628.

5. Ridgeon EE, Young PJ, Bellomo R, Mucchetti M, Lembo R, Landoni G. The Fragility Index in Multicenter Randomized Controlled Critical Care Trials. Crit Care Med 2016;44: 1278–1284.

6. Docherty KF, Campbell RT, Jhund PS, Petrie MC, McMurray JJ. How robust are clinical trials in heart failure? Eur Heart J 2017;38: 338–345.

7. Ahmed W, Fowler RA, McCredie VA. Does Sample Size Matter When Interpreting the Fragility Index? Crit Care Med 2016;44: e1142–e1143.

8. R Development Core Team. R: A Language and Environment for Statistical Computing. Vienna, Austria: R Foundation for Statistical Computing, 2016.

9. Therneau TM and Grambsch PM. Modeling survival data: Extending the Cox Model. New York: Springer, 2000. ISBN: 0-387-98784-3.

10. Cancer Genome Atlas N. Comprehensive molecular characterization of human colon and rectal cancer. Nature 2012;487: 330–337.

11. Lichman M. UCI Machine Learning Repository. Irvine, CA, 2013.

12. Carter RE, McKie PM, Storlie CB. The Fragility Index: a P value in sheep’s clothing? European Heart Journal 2017; 38: 346–348.

