## Supplementary Material for "fragilityindex: An R Package for Statistical Fragility Estimates in Biomedicine"

**Supplementary Materials**

*Fragility Index for contingency tables*

Below is the complete R code to generate the final fragility indices from the clinical data as described in the main manuscript.

```
### Preliminaries
library(data.table)
library(fragilityindex)
options(stringsAsFactors = FALSE)

### Read in clinical study master file
### This is the no-clob file nolf file created as per the
### README_R.docx file included in the AACT downloaded directory
# Primary key: NCT_ID
cs <- read.delim("AACT201603_pipe_delimited/clinical_study_noclob_nolf.txt",
                 sep="|",
                 quote="",
                 na.strings = "",
                 comment.char="",
                 encoding="UTF-8",
                 header=TRUE)

### Read in results_outcomes file
# Primary key: OUTCOME_ID
ro <- read.delim("AACT201603_pipe_delimited/results_outcomes_nolf.txt",
                 sep="|",
                 quote="",
                 na.strings = "",
                 comment.char="",
                 encoding="UTF-8",
                 header=TRUE)

### Read in results_outcomes_measure file
# Primary key: OUTCOME_ID
rom <- read.delim("AACT201603_pipe_delimited/results_outcome_measure_nolf.txt",
                  sep="|",
                  quote="",
                  na.strings = "",
                  comment.char="",
                  encoding="UTF-8",
                  header=TRUE)

### Read in results_outcomes_analysis file
# Primary key: OUTCOME_ID
roa <- read.delim("AACT201603_pipe_delimited/results_outcome_analysis_nolf.txt",
                  sep="|",
                  quote="",
                  na.strings = "",
                  comment.char="",
                  encoding="UTF-8",
                  header=TRUE)

# Get only clinical trials which used chi-square or fisher exact tests as their
# reported method of analysis of a given outcome
roa.dt <- as.data.table(roa)
roa.fechi <- roa.dt[METHOD %in% c("Fisher Exact")] # fisher only

### Read in results_outcome_measure file
# Primary key: OUTCOME_ID
romc <- read.delim("AACT201603_pipe_delimited/results_outcome_measure_ctgy_nolf.txt",
                   sep="|",
                   quote="",
                   na.strings = "",
                   comment.char="",
                   encoding="UTF-8",
                   header=TRUE)

# Join results_outcome_measure to results_outcome_measure_category
romc.sub <- romc[,c("OUTCOME_MEASURE_ID","OUTCOME_MEASURE_CATGY_ID","CATEGORY_TITLE","OUTCOME_VALUE")]
rom.sub <- rom[,c("OUTCOME_ID","OUTCOME_MEASURE_ID","UNIT_OF_MEASURE","MEASURE_TYPE")]
rom.romc <- merge(rom.sub, romc.sub, by.x="OUTCOME_MEASURE_ID", by.y="OUTCOME_MEASURE_ID")

# Join rom.romc to results_outcome table
ro.sub <- ro[,c("NCT_ID","OUTCOME_ID","OUTCOME_TYPE")]
ro.rom.romc <- merge(ro.sub, rom.romc, by.x="OUTCOME_ID", by.y="OUTCOME_ID")

# Join ro.rom.romc to results_outcome_analysis table
roa.fechi <- as.data.frame(roa.fechi)
roa.sub <- roa.fechi[,c("OUTCOME_ID","RESULTS_OUTCOME_ANALYSIS_ID","PARAM_TYPE","NON_INFERIORITY","P_VALUE","METHOD")]
ro.rom.romc.roma <- merge(ro.rom.romc, roa.sub, by.x="OUTCOME_ID", by.y="OUTCOME_ID")
ro.rom.romc.roma$CATEGORY_TITLE <- NULL

d1 <- as.data.table(ro.rom.romc.roma)
sort(table(d1[,length(RESULTS_OUTCOME_ANALYSIS_ID), by=.(OUTCOME_ID)]$V1))
d1[,logicalValue:=ifelse(length(RESULTS_OUTCOME_ANALYSIS_ID)==4,1,0), by=.(OUTCOME_ID)]
d1.4 <- d1[logicalValue==1]
d1.4[,logicalValue:=NULL]

### Fix p values, which were badly formatted strings
trim.leading <- function (x){sub("^\\s+", "", x)}
trim.trailing <- function (x){sub("\\s+$", "", x)}
pvals <- gsub(">", "", d1.4$P_VALUE)
pvals <- gsub("<", "", pvals)
pvals <- gsub("=", "", pvals)
pvals <- trim.leading(pvals)
pvals <- trim.trailing(pvals)
pvals <- as.numeric(pvals)
d1.4$pvals <- pvals
d1.4s <- d1.4[pvals<0.05]

# omit outcome_id 78749 which has malformed values
d1.4s <- d1.4s[OUTCOME_ID!=78749]
d1.4s$OUTCOME_VALUE <- as.numeric(d1.4s$OUTCOME_VALUE)
d1.4s[, n:=ifelse(OUTCOME_VALUE < max(OUTCOME_VALUE),"n_event","n_total"), by=.(OUTCOME_ID, OUTCOME_MEASURE_ID)]

### fragility.index calculations
uniq_ncts <- unique(d1.4s$NCT_ID)

out.df <- as.data.frame(matrix(NA, nrow=length(uniq_ncts), ncol=6))
d1.4s$P_VALUE <- NULL # delete useless column
d1.4s$NON_INFERIORITY <- NULL # delete useless column
setkey(d1.4s, NCT_ID)

for(i in 1:length(uniq_ncts)){
    nct <- uniq_ncts[i]

    temp.dt <- d1.4s[NCT_ID==nct]
    print(nct)

    outcome_ids <- unique(temp.dt$OUTCOME_MEASURE_ID)

    nevent1 <- temp.dt[OUTCOME_MEASURE_ID==outcome_ids[1] & n=="n_event"]$OUTCOME_VALUE
    ntotal1 <- temp.dt[OUTCOME_MEASURE_ID==outcome_ids[1] & n=="n_total"]$OUTCOME_VALUE
    nevent2 <- temp.dt[OUTCOME_MEASURE_ID==outcome_ids[2] & n=="n_event"]$OUTCOME_VALUE
    ntotal2 <- temp.dt[OUTCOME_MEASURE_ID==outcome_ids[2] & n=="n_total"]$OUTCOME_VALUE

    cat(paste(i, ":  ", nevent1, ntotal1, nevent1, ntotal1, "\n"))

    findex <- tryCatch(fragility.index(nevent1, nevent2, ntotal1, ntotal2), error=function(e){return(NA)} )
    findex <- unlist(findex)

    try(out.df[i,1] <- nct)
    try(out.df[i,2] <- nevent1)
    try(out.df[i,3] <- ntotal1)
    try(out.df[i,4] <- nevent2)
    try(out.df[i,5] <- ntotal2)
    try(out.df[i,6] <- findex)
}

names(out.df) <- c("NCT","nevent1","ntotal1","nevent2","ntotal2","fragility.index")
out.df2 <- na.omit(out.df) # final data frame
```

*Fragility Index for survival analysis*

The R code below can perform the fragility index survival analysis on the TCGA data described in the main manuscript.

```
set.seed(1000) # To ensure reproducible results

### Preliminaries
rm(list=ls())
library(fragilityindex)
library(data.table)
library(stringr)
library(survival)

# Read in and prepare Copy Number Alteration TCGA table
in2 <- fread("CNA_Genes.txt")

in.sub <- in2[Gene %in% c("BCL2L1","POFUT1"),]

bcl.cases <- in2[Gene=="BCL2L1",Cases]
pofut.cases <- in2[Gene=="POFUT1",Cases]

bcl.case0 <- unlist(str_split(bcl.cases,","))
pofut.case0 <- unlist(str_split(pofut.cases,","))

bcl.case1 <- str_sub(bcl.case0, end = -4L)
pofut.case1 <- str_sub(pofut.case0, end = -4L)

# Read in and prepare survival data/clinical TCGA information
in1 <- fread("data_clinical.txt")

in1.sub1 <- in1[PATIENT_ID %in% c(bcl.case1, pofut.case1)]
in1.sub2 <- in1[!(PATIENT_ID %in% c(bcl.case1, pofut.case1))]

in1.sub1[,strat:="mutated"]
in1.sub2[,strat:="not_mutated"]

in1.sub1$OS_MONTHS[which(in1.sub1$OS_MONTHS==0)] <- max(in1.sub1$OS_MONTHS)
in1.sub2$OS_MONTHS[which(in1.sub2$OS_MONTHS==0)] <- max(in1.sub2$OS_MONTHS)

in1.sub1$months <- in1.sub1$OS_MONTHS
in1.sub2$months <- in1.sub2$OS_MONTHS

in1.com <- rbind(in1.sub1, in1.sub2)
in1.com$status <- as.factor(in1.com$OS_STATUS)
levels(in1.com$status) <- c(1,0)
in1.com$status <- as.numeric(as.character(in1.com$status))
in1.com$strat <- as.factor(in1.com$strat)

com <- as.data.frame(in1.com)
com <- com[,c("months","status","strat")]
com2 <- na.omit(com)

### Plot survival curve
surv.obj <- survfit(Surv(months, status) ~ strat, data=com2)
plot(surv.obj)

### Survival curve statistics
surv.diff <- survdiff(Surv(months, status) ~ strat, data=com2)
print(s1)

### Run the fragility analysis
# Takes about 1 minute for 1000 iterations on 1.7 Ghz Skylake i7 processor (Macbook Air)
survivalfragility(Surv(months, status) ~ strat, data=com2, niter=1000, progress.bar = TRUE)
```

*Fragility Index for logistic regression β-coefficients*

The following R code can be used to perform fragility index calculation for logistic regression β-coefficients on the example from UCI Machine Learning Repository described in the main text.

> library(car)

>mydata=read.csv("http://archive.ics.uci.edu/ml/machine-learning-databases/heart-disease/processed.cleveland.data")

>colnames(mydata) =c("age","sex","cp","trestbps","chol","fbs","restecg","thalach",

"exang","oldpeak","slope","ca","thal","num")

> mydata$ca=as.numeric(mydata$ca)

> mydata$thal=as.numeric(mydata$thal)

The “num” variable reflects heart disease status and is originally in a severity scale (1-4). For the current analysis, we have to transform the outcome into a binary 0/1 outcome (all values above 0 become 1).

> mydata$num <- recode(dat$num, "1:4=1")

With the data properly formatted, we first should determine whether any covariates in the model are collinear. In the current analysis, we use variance inflation factor (VIF) to measure multicolinearity although there are many other methods.

```
> vif(glm(num ~ age + sex +cp + trestbps + chol + fbs + restecg + thalach + exang + oldpeak + slope + ca + thal, family="binomial",data = mydata))

# output is displayed in table format for readability

	 VIF
age      1.493313
sex      1.303031
cp 	 1.316989
trestbps 1.207288
chol     1.136903
fbs 	 1.073972
restecg  1.089937
thalach 	 1.646313
exang  	 1.376995
oldpeak  1.753450
slope    1.670978
ca       1.354936
thal	 1.520322
```

As all covariate VIF values are below 5 (this value is not a hard requirement but has been suggested to use as a lower indicating threshold), we can conclude that there is not a strong correlation between any coefficients. Next, we can perform an initial, basic logistic regression for preliminary evaluation and to better understand the fragility index outcomes for each coefficient.

```
> lmfit <- glm(num~age + sex +cp + trestbps + chol + fbs + restecg + thalach + exang + oldpeak + slope +ca +thal,family="binomial",data=mydata)
> summary(lmfit)

Call:
glm(formula = num ~ age + sex + cp + trestbps + chol + fbs +
    restecg + thalach + exang + oldpeak + slope + ca + thal,
    family = "binomial", data = mydata)

Deviance Residuals:
    Min       1Q   Median       3Q      Max
-2.7767  -0.5246  -0.1861   0.4247   2.3695

Coefficients:
             Estimate Std. Error z value Pr(>|z|)
(Intercept) -7.414600   2.879054  -2.575 0.010014 *
age         -0.012937   0.024001  -0.539 0.589873
sex          1.312155   0.487786   2.690 0.007145 **
cp           0.560374   0.192026   2.918 0.003520 **
trestbps     0.023857   0.010726   2.224 0.026128 *
chol         0.004900   0.003766   1.301 0.193300
fbs         -0.956923   0.560674  -1.707 0.087871 .
restecg      0.252305   0.185128   1.363 0.172923
thalach     -0.020484   0.010199  -2.009 0.044588 *
exang        0.913115   0.413365   2.209 0.027176 *
oldpeak      0.252390   0.211818   1.192 0.233441
slope        0.592078   0.363295   1.630 0.103155
ca           1.250928   0.265662   4.709 2.49e-06 ***
thal         0.346178   0.100237   3.454 0.000553 ***
---
Signif. codes:  0 ‘***’ 0.001 ‘**’ 0.01 ‘*’ 0.05 ‘.’ 0.1 ‘ ’ 1

(Dispersion parameter for binomial family taken to be 1)

    Null deviance: 408.71  on 295  degrees of freedom
Residual deviance: 204.03  on 282  degrees of freedom
  (6 observations deleted due to missingness)
AIC: 232.03

Number of Fisher Scoring iterations: 6
```

These results signify that there are a couple of predictor variables that are significantly associated with heart disease risk. By next performing the “logisticfragility” function, we can determine the fragility index for all significant variables noted above.

```
> logisticfragility(num ~ age + sex +cp + trestbps + chol + fbs + restecg + thalach + exang + oldpeak + slope + ca + thal, data = mydata, covariate="all", niter=100, progress.bar=FALSE)

   coefficient fragility.index
1  (Intercept)           15.92
2          age            0.00
3          sex           33.10
4           cp           35.10
5     trestbps           10.45
6         chol            0.00
7          fbs            0.00
8      restecg            0.00
9      thalach            5.99
10       exang           12.67
11     oldpeak            0.00
12       slope            0.00
13          ca           68.84
14        thal           64.00
```

The first thing to notice is that only variables with significant associations in the model will have a fragility index calculated for them.
